# Precise Alterations in Hippocampal Neural Deterioration and Neural Correlates of Visual Short-Term Memory in Individuals with Amnestic Mild Cognitive Impairment

**DOI:** 10.1101/2023.09.21.558904

**Authors:** Ye Xie, Tinghao Zhao, Yunxia Li, Yixuan Ku

## Abstract

**Purpose:** Atrophy of the hippocampus is an early biomarker of Alzheimer’s disease (AD) and a main contributor to the patients’ mnemonic degeneration. While previous research has mostly focused on how hippocampal atrophy impaired long-term memory performance, its relation to short-term memory impairment, which was also found among patients with AD, remains largely uninvestigated. Filling this gap, the current study examined how atrophy in the hippocampus and its subfields may have influenced visual short-term memory (VSTM) in patients with amnestic mild cognitive impairment (aMCI), a common precursor of AD.

**Methods:** Fifty-eight aMCI patients and 69 healthy controls (HC) matched in age were included in the current study. VSTM was assessed using an adapted change detection task with a memory load of 2 or 4 items. Hippocampal subfields were automatically segmented in T1-weighted image using FreeSurfer and were manually inspected for errors. Volumes of the subfields were extracted and compared between aMCI and HC subjects using ANCOVA with age, gender and education as covariates. Furthermore, we also examined the partial correlation between VSTM performances and hippocampal subfield volumes with age, gender and education as covariates.

**Results:** Compared to HC subjects, aMCI subjects had lower response accuracy (ACC) and lower memory capacity under both load conditions and had longer reaction time (RT) in the 2-load condition. Left hippocampus volume was significantly smaller in aMCI and was positively correlated with ACC and capacity in HC but not in aMCI. Among the hippocampal subfields, left hippocampal tail, left molecular layer, left dentate gyrus (DG), left CA4, bilateral subiculum, bilateral presubiculum, bilateral fimbria, were significantly smaller while right hippocampal fissure was significantly widened in aMCI compared to HC. Volumes of the left subiculum, left molecular layer, left DG and left CA4 were positively correlated with ACC and capacity in HC but not in aMCI. Bilateral fimbria volume was negatively correlated with RT under the 2-load condition in HC but not in aMCI.

**Conclusion:** The results of this study suggested that hippocampal deterioration, especially in subfields related to information input and output (e.g. molecular layer, DG, subiculum), may have contributed to VSTM impairment in aMCI by disrupting hippocampal-cortical communications. This finding adds to increasing evidence of hippocampal engagement in short-term memory processes and points to VSTM impairment as a potential neuropsychological indicator for MCI and AD.

## Introduction

Alzheimer’s disease (AD) is a progressive neurodegenerative disorder which causes severe cognitive decline in elderly people. Amnestic Mild Cognitive Impairment (aMCI) is a transitional stage between normal age-related cognitive decline and abnormal cognitive impairment caused by AD (Li & Zhang, 2015; Petersen et al., 2006). Individuals diagnosed with aMCI often experience memory problems greater than what is expected for their age but do not meet the criteria for dementia. While aMCI can affect various cognitive domains, one of the most prominent and early signs is in the domain of visual short-term memory (VSTM), which has been recently flagged as early marker of disease (Norton et al., 2020; Parra et al., 2022). VSTM plays a fundamental role in our everyday lives (A. D. Baddeley et al., 2011). It is responsible for temporarily holding and manipulating visual information, allowing us to perform a wide range of cognitive tasks, such as recognizing faces, reading, and navigating our environment. Literatures have consistently reported deficits in VSTM not only in MCI or AD, but also asymptomatic familial AD who still exhibited normal cognitive function (Kirova et al., 2015; Parra et al., 2010; Parra et al., 2011; Zokaei et al., 2020). These deficits manifest as reduced accuracy and increased susceptibility to interference when holding visual information for brief periods. Understanding the neural underpinnings of these VSTM deficits is essential for both diagnosis and intervention. However, the underlying pathology of the deficits in VTSM in aMCI individuals remained unclear.

The typical pathology of AD has been characterized by the brain atrophy spreading from hippocampus or medial temporal lobe (MTL) to the rest of brain areas (Thompson et al., 2007). The atrophy of hippocampus has been widely reported in the case of MCI and AD and considered as AD pathological hallmarks. The degree of the hippocampal atrophy has been proven to be not only predictive of progression from MCI to AD but also useful in distinguishing AD from normal elderly people (Scheltens et al., 1992; Visser et al., 2002). Hippocampus, the seahorse-shaped structure nestled deep within the brain’s medial temporal lobe, has long been recognized as a critical player in memory processes ever since the 1957 report of the case study H.M. (A. Baddeley et al., 2011; Squire, 2009; von Allmen et al., 2013). Hippocampus has been traditionally regarded as the core area engaged in episodic memory, which facilitated multiple studies investigating the association between hippocampus atrophy and episodic memory in aMCI. Recently, the critical role of hippocampus in VSTM has been revealed by several studies. During the VSTM tasks, hippocampus has been found to be firing specific neuronal to maintain the representations of items and the wellness of the maintain has been found to correlated with the hippocampus theta activity (Boran et al., 2022; Liu et al., 2020). Furthermore, we also observed the dissociation between VSTM performance and the atrophic hippocampus among the aMCI individuals. All these findings suggested the association between the dysfunction of hippocampus and the deficits in VSTM in the aMCI.

Notably, hippocampus has intricate and heterogeneous structure and can be divided into subfields with different functional characteristics varied along the anterior-posterior axis. For example, it has been suggested that CA2/3 and dentate gyrus are input structures which are responsible for encoding when considering their role in memory, and CA1 and subiculum are output structures responsible for retrieval (Eldridge et al., 2005; Nauer et al., 2015; Zammit et al., 2017; Zeineh et al., 2003). Imaging studies have revealed heterogeneity of these subfields when hippocampus is affected by AD. CA1, subiculum and dentate gyrus were the most frequently reported to be found atrophic when compared the AD or MCI patients to normal controls and has been reported to be related with the deficits in mnemonic processing (Apostolova et al., 2006; Apostolova, Mosconi, et al., 2010; Broadhouse et al., 2019; de Flores et al., 2015). Subfields analysis has suggested that the prevalent atrophy of the presubicular-subicular complex appears early during the progression of AD and presubiculum and subiculum could be considered as possible predictors of verbal memory performance in MCI (Carlesimo et al., 2015). However, little is known about how is the deterioration of inner hippocampus would affect the deficit of VSTM in aMCI individuals.

Accordingly, in the current study, we will delve into the precise alterations in hippocampal neural deterioration and exploring the neural correlates of VSTM deficits in individuals with aMCI. By elucidating the intricate relationship between hippocampal deterioration and VSTM deficits, we aim to contribute to the growing body of knowledge surrounding aMCI and provide valuable insights into potential avenues for therapeutic intervention and cognitive rehabilitation.

## Method

### 2.1 Diagnostic assessment

The participants in the current study were recruited from Tongji Hospital in Shanghai, China. aMCI was designated by a neurologist from the department of Neurology and the Department of Memory Clinic of Shanghai Tongji Hospital based on the ‘MCI-criteria for the clinical and cognitive syndrome’(Albert et al., 2011). Specifically, the criteria included: (a) The participants and their caregiver had complaint of memory/cognitive decline; (b) Mini-Mental State Examination (MMSE) or MoCA scores met criteria adjusted by education; (c) Clinical Dementia Rating Scale (CDR) of 0.5; (d) met any of the following three criteria: (1) at least two performances within a cognitive domain fell below the established cutoff (> 1SD); (2) at least two cognitive domains were impaired (> 1SD); (3) had more than one function described in instrumental activities of daily living (IADL-14) scored of 1 or more.

### 2.2 Participants

The current study included fifty-eight aMCI subjects and sixty-nine normal elderly people (NC) (For demographic characteristic, please see Supplementary Table 1). Informed consent was obtained from all participants and the study was approved byIRB board of Tongji Hospital with ethical and safety guidelines consistent with the Declaration of Helsinki.

### 2.3 VSTM task

VSTM was measured by an adopted change detection task (Fig. 1) (Vogel & Machizawa, 2004). Participants were asked to indentify whether the test array was either identical to the memory array or differed by one color. The presentation of test array would last until participants gave their response. Accuracy (ACC) and mean reaction time (RT) were calculated for each participants, and capacity was represented by Cowan’s K coefficient (load x (hit rate -false alarm rate)) (Cowan, 2001).

**Figure 1.**
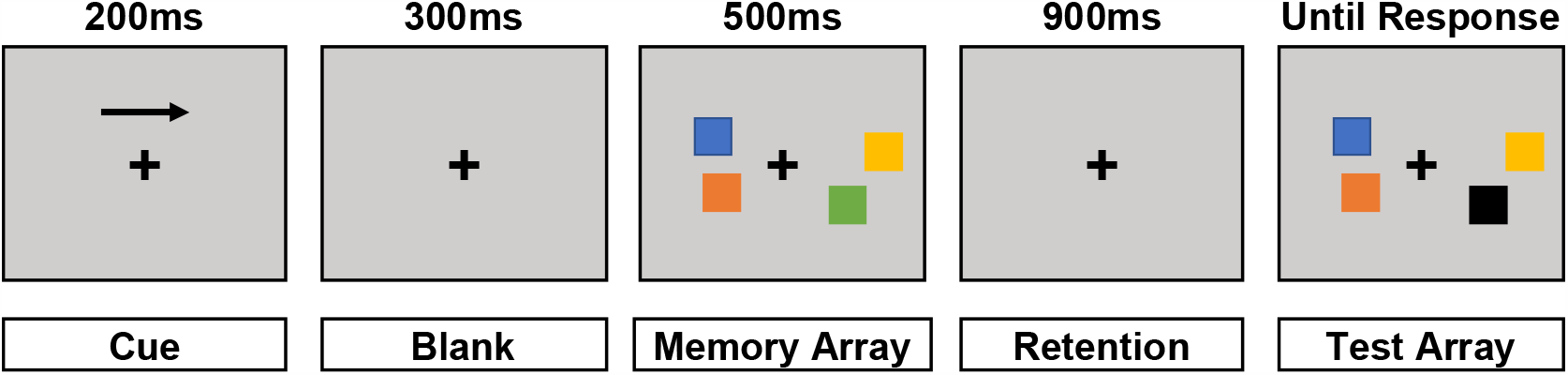
Stimuli and trail procedure of visual change detection task.

### 2.4 MRI Acquisition

Image data were collected from two sites: East China Normal University with Siemens Prisma machine (81 participants, 38 aMCI participants and 43 NC participants) and Tongji Hospital using Siemens Verio machine (46 participants, 20 aMCI participants and 26 NC participants). To delineate the influence of machine, collecting site would be use as covariate in imaging analysis.

Collecting parameters of these two sites were same in TE (2.98ms), TR (2530ms), FOV (256mm x 256mm), and acquisition matrix (256 x 256), but different in the reconstruction voxel size (for East China Normal University: 0.5 x 0.5 x 1.0 mm^3^. For Tongji Hospital: 1.0 x 1.0 x 1.0mm^3)^.

Then the segmentation of hippocampal subfield was conducted based a delicate template of hippocampal subfileds with freesurfer software. We focused on the parasubiculum, presubiculum, subiculum, CA1, CA3 (or CA2/3), CA4, dentate gyrus (DG), hippocampal-amygdaloid transition region (HATA), fimbria, molecular layer, hippocampal fissure and hippocampal tail. After finishing the segmentation, the goodness of the segmentation was manually checked and those with bad segmentation were included from the study.

### 2.5 Statistical methods

Repeated measures ANOVA was used to test the ACC and the RT of the VSTM tasks ad one-way analysis of covariance (ANCOVA) was used to compared participants’ capacity of VSTM. Age, gender, education and collecting site were used as covariates and Bonferroni method was applied for multiple comparison correction. One-way analyses of covariance (ANCOVAs) were also used for comparison of gray matter volumes of hippocampal subfields with age, gender, education, site and TIV as covariates, and FDR method were used for multiple comparison correction. Partial correlation was used to test the correlation between the volume of hippocampal subfields showing significant between-group difference and performance of VSTM task with age, gender, education, site and TIV as covariates.

## Result

### 3.1 demographic characteristic of included sample

Twelve subfields were constructed (Fig. 2). Three subjects were excluded from the analysis because of the poor segmentation of hippocampal subfields, remaining 54 aMCI and 66 NC. demographic characteristics were shown in the Table 1.

**Table 1.** demographic characteristics.

|  | aMCI<br>(n = 54) | NC<br>(n = 66) | Between-group differences |
| --- | --- | --- | --- |
| Age (years) | 72.42 (6.75) | 70.35 (6.12) | $t(118) = 1.77, p = 0.080^a$ |
| Gender (male: female) | 23:31 | 43:23 | $\chi^2 = 6.11, p = 0.013^b$ |
| Education (years) | 11.37 (2.66) | 12.42 (2.80) | $t(118) = -2.1, p = 0.038^a$ |
| MMSE | 24.93 (2.49) | 27.49 (2.51) | $W = 621.00, p < 0.001^c$ |
| MoCA | 17.17 (3.50) | 24.24 (2.51) | $W = 183.00, p < 0.001^c$ |

Values are mean (standard deviation). Abbreviation: aMCI, amnesic mild cognitive impairment; NC, normal controls; MMSE, Mini-Mental State Examination; MoCA, Montreal Cognitive Assessment.
<sup>a</sup> independent two-sample t-test, <sup>b</sup> chi-square test, <sup>c</sup> Mann-Whitney U test.

**Figure 2.**
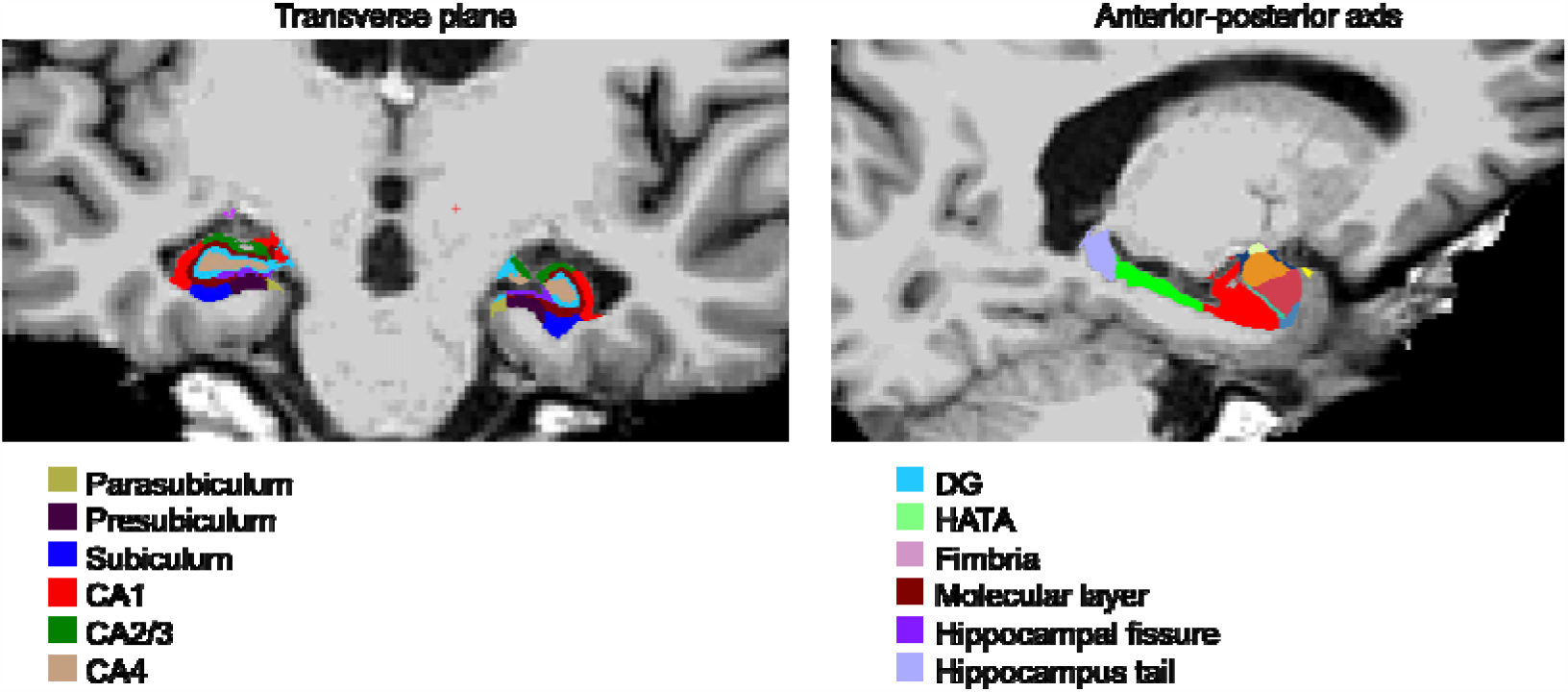
Segmentation of hippocampal subfields. Abbreviation: DG, dentate gyrus; HATA, hippocampal-amygdaloid transition region.

### 3.2 VSTM performance

ANCOVA analysis showed significant main effect of group for ACC (F (1, 114) = 15.28, *p* < 0.001) but no interaction between group and task load (*p* = 0.21) after covarying the effect of age, gender, education and collecting site. *Post-hoc* analysis with Bonferroni correction showed that aMCI group performed significant worse than the NC group in the task of 2-item load (aMCI: 0.88±0.11; NC: 0.95±0.051; *p* < 0.001) and marginally significant worse in the task of 4-item load (aMCI: 0.73±0.095; NC: 0.78±0.091; *p* = 0.051) with age, gender, education and collecting site as covariates.

Significant main effect of group (*F(1,114) = 5*.*119, p* = 0.026) and interaction between group and task load (F (1, 114) = 9.78, *p* = 0.002) was found for RT after covarying the effect of age, gender, education and collecting site. *Post-hoc* analysis with Bonferroni correction showed significant longer RT in aMCI group in task of 2-item load (*p = 0*.*009*), but not in the task of 4-item load with age gender, education and collecting site as covariates.

Significant between-group difference was found in capacity (F (1, 114) = 8.46, *p* < 0.004; aMCI: 1.93±0.11; NC: 2.40±0.089).

3.2 Between-group difference in hippocampal volume and correlation to VSTM performance

Structural comparisons were conducted for bilateral hippocampus and all the subfield by ANCOVA, with age, gender, education, collecting site and TIV as covariates. The results were then corrected by FDR method. Results showed that significant smaller gray matter volume was found in the aMCI group in the left lateral hippocampus (Figure 3A), and among the hippocampal subfields, left hippocampal tail, left molecular layer, left DG, left CA4, bilateral subiculum, bilateral presubiculum, and bilateral fimbria were found significantly smaller in the aMCI group when compared to NC while right hippocampal fissure showed right hippocampal fissure in the aMCI group (Figure 3B and Table 2). No main effect of group was found for right lateral hippocampus or other subfields (*ps* > 0.05).

**Table 2.** Subfields showing significant between-group difference.

| subfield | group | mean (SD) | ANCOVA |
| --- | --- | --- | --- |
| L hippocampus tail | aMCI | 450.73(12.02) | F(1,113) = 5.71<br>$p = 0.039$ |
|  | NC | 509.16(8.9) |  |
| L subiculum | aMCI | 380.81(8.34) | F(1,113) = 6.56<br>$p = 0.039$ |
|  | NC | 425.33(6.24) |  |
| L presubiculum | aMCI | 254.06(6.52) | F(1,113) = 5.94<br>$p = 0.039$ |
|  | NC | 287.15(5.48) |  |
| l molecular | aMCI | 463.06(10.13) | F(1,113) = 8.79<br>$p = 0.026$ |
|  | NC | 519.28(6.68) |  |
| L DG | aMCI | 240.31(5.31) | F(1,113) = 9.93<br>$p = 0.026$ |
|  | NC | 272.5(3.9) |  |
| L CA4 | aMCI | 212.27(4.44) | F(1,113) = 7.06<br>$p = 0.037$ |
|  | NC | 237.28(3.36) |  |
| L fimbria | aMCI | 64.86(3.61) | F(1,113) = 6.78<br>$p = 0.037$ |
|  | NC | 84.56(2.88) |  |
| R subiculum | aMCI | 391.17(8.87) | F(1,113) = 6.11<br>$p = 0.039$ |
|  | NC | 434.38(6.1) |  |
| R hippocampus fissure | aMCI | 184.88(5.49) | F(1,113) = 5.81<br>$p = 0.039$ |
|  | NC | 178.79(5.74) |  |
| R presubiculum | aMCI | 242.96(5.89) | F(1,113) = 8.55<br>$p = 0.026$ |
|  | NC | 274.97(5.17) |  |
| R fimbria | aMCI | 59.77(3.2) | F(1,113) = 7.09<br>$p = 0.037$ |
|  | NC | 73.09(2.87) |  |
| L hippocampus | aMCI | 2888.41(61.09) | F(1,113) = 9.70<br>$p = 0.026$ |
|  | NC | 3244.81(40.96) |  |

P-values were FDR corrected. Abbreviation: DG, dentate gyrus.

**Figure 2.**
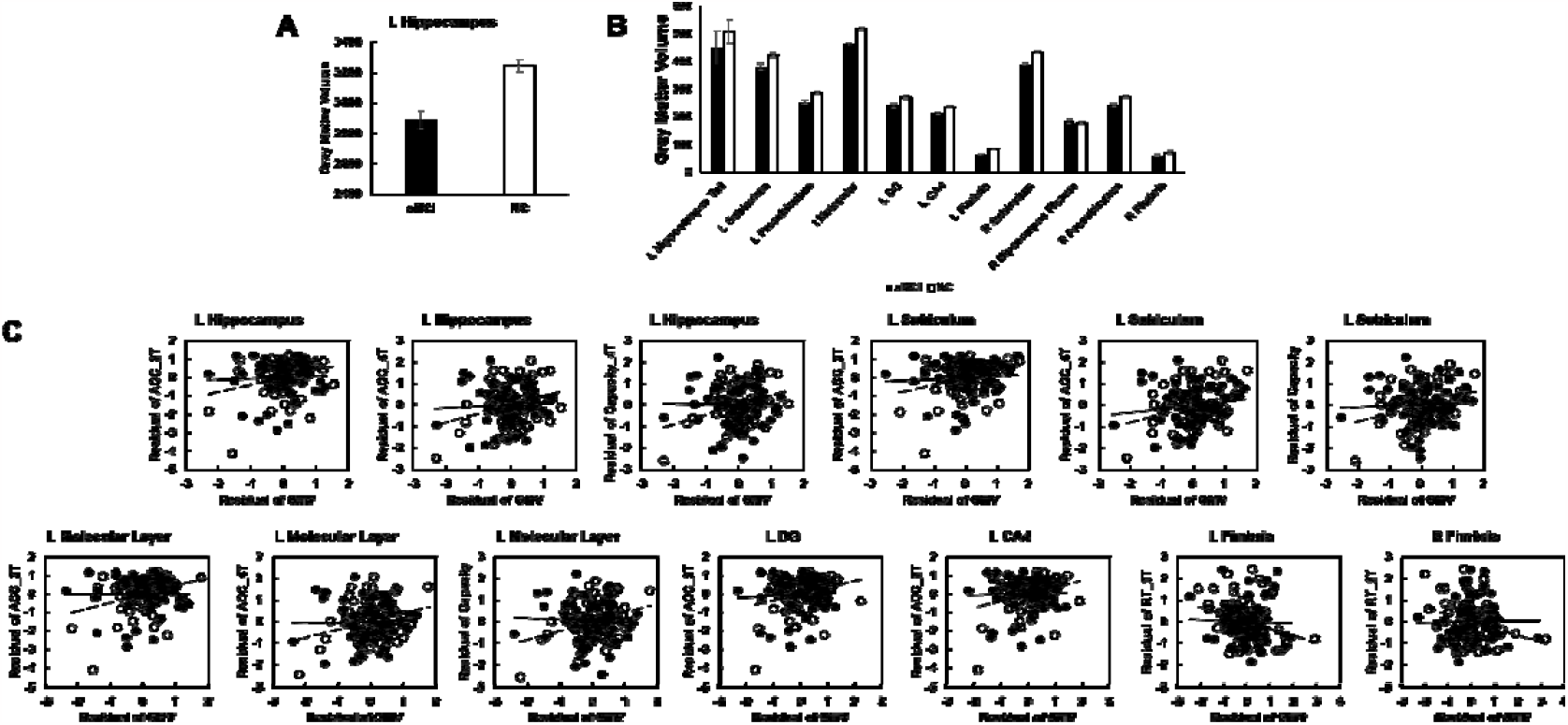
Significant between-group difference in unilateral hippocampus (A) and hippocampal subfields (B). Correlation between gray matter volume (GMV) and VSTM performance (C) were conducted by partial correlation analysis. Analyses were conducted with age, gender, education, TIV and collecting site as covariates. Abbreviation: DG, dentate gyrus.

Partial correlation analysis with age, gender, education, collecting site and TIV as covariates showed that the gray matter volume of the left hippocampus, left subiculum, left molecular, left DG and left CA4 were correlated with the accuracy or capacity, and bilateral fimbria were correlated to the RT of VSTM tasks (*ps* < 0.05, Figure 2C and Table 3) in the NC group but no correlation was found in the aMCI group (*ps* > 0.05).

**Table 3.** Brain-behavioral association between hippocampal subfield and VSTM performance.

| subfield | Group | ACC_2T | RT_2T | Capacity_2T | ACC_4T | RT_4T | Capacity_4T |
| --- | --- | --- | --- | --- | --- | --- | --- |
| L subiculum | aMCI | \ | \ | \ | \ | \ | \ |
| | NC | $r = 0.297$,<br>$p = 0.020$ | \ | \ | $r = 0.317$,<br>$p = 0.013$ | \ | $r = 0.301$,<br>$p = 0.018$ |
| L molecular layer | aMCI | \ | \ | \ | \ | \ | \ |
| | NC | $r = 0.328$,<br>$p = 0.010$ | \ | \ | $r = 0.280$,<br>$p = 0.029$ | \ | $r = 0.309$,<br>$p = 0.015$ |
| L DG | aMCI | \ | \ | \ | \ | \ | \ |
| | NC | $r = 0.270$,<br>$p = 0.035$ | \ | \ | \ | \ | \ |
| L CA4 | aMCI | \ | \ | \ | \ | \ | \ |
| | NC | $r = 0.294$,<br>$p = 0.022$ | \ | \ | \ | \ | \ |
| L fimbria | aMCI | \ | \ | \ | \ | \ | \ |
| | NC | \ | $r = -0.288$,<br>$p = 0.024$ | \ | \ | \ | \ |
| r fimbria | aMCI | \ | \ | \ | \ | \ | \ |
| | NC | \ | $r = -0.283$,<br>$p = 0.027$ | \ | \ | \ | \ |
| L hippocampus | aMCI | \ | \ | \ | \ | \ | \ |
| | NC | $r = 0.282$,<br>$p = 0.027$ | \ | \ | $r = 0.294$,<br>$p = 0.021$ | \ | $r = 0.323$,<br>$p = 0.011$ |

\ represents insignificant correlation. Abbreviation: DG, dentate gyrus.

## Discussion

In the current study, we conducted structural comparison in hippocampal subfield to target precise atrophy of hippocampus in individual with aMCI and identified the association between the atrophy and VSTM deficits. The current results showed that compared to HC subjects, aMCI subjects had lower response accuracy (ACC) and lower memory capacity under both load conditions and had longer reaction time (RT) in the 2-load condition. Left hippocampus volume was significantly smaller in aMCI and was positively correlated with ACC and capacity in HC but not in aMCI. Among the hippocampal subfields, left hippocampal tail, left molecular layer, left DG, left CA4, bilateral subiculum, bilateral presubiculum and bilateral fimbria, were significantly smaller and right hippocampal fissure were significant widened in aMCI compared to HC. Volumes of the left subiculum, left molecular layer, left DG and left CA4 were positively correlated with ACC and capacity in HC but not in aMCI. Bilateral fimbria volumes were negatively correlated with RT under the 2-load condition in HC but not in aMCI.

Consistent with previous findings, our results revealed the VSTM deficits. The aMCI groups not only exhibited lower accuracy and slower response but also weaker processing capacity. VSTM has been flagged as cognitive indicator for early AD recently (Alescio-Lautier et al., 2007; Kessels et al., 2015; Zokaei & Husain, 2019; Zokaei et al., 2020). Studies have reported the association between VSTM performance and early metabolism pathology of AD and highlighted the predictive role of VSTM to distinguish MCI of early stage from normal controls (Norton et al., 2020; Parra et al., 2022). Hippocampus, a brain area traditionally considered as core area for long-term memory, has been revealed to be engaged and play important role in VSTM processing (Boran et al., 2022; Liu et al., 2020). In the current study, the gray matter volume of left hippocampus was significant smaller in the aMCI group than in the controls, convergent with the previous studies showing left lateral hippocampus as the most vulnerable brain regions affected by AD pathology (Ferreira et al., 2011). The degree of hippocampal atrophy has been regarded as a hallmark of AD progression (Scheltens et al., 1992; Visser et al., 2002. Our results also revealed that the left lateral hippocampus was correlated with the VSTM performance in the controls but not in the aMCI group. Such results add evidence to the notion that hippocampus support VSTM. In contrast, individuals with aMCI suffer from hippocampal atrophy, and thus, exhibited disrupted function of hippocampus in supporting VSTM.

When look into the subdivision of hippocampus, left hippocampus tail, left molecular layer, left DG, left CA4, bilateral fimbria, bilateral subiculum and bilateral presubiculum showed significant atrophy in the aMCI group when compared to the NC group, while right hippocampal fissure showed dilatation in the aMCI group. Hippocampus has an intricate structure and has been divided into subfields according to their distinct functional characteristics along the anterior-posterior axis. The atrophic subfields found in the current study might reveal that the deterioration is heterogeneous inner hippocampus along the AD progression.

Most of the subfields showing between-group differences were subdivisions suggested as pathway for information input or output in hippocampus. Molecular layer was the input to dentate gyrus (Forster et al., 2006). Dentate gyrus combined the information conveyed by entorhinal cortex and associational projections arising from this area has been regarded to be specifically organized to integrate distant levels of the hippocampal formation (Amaral & Witter, 1989; Braak et al., 1996; Knierim, 2015). It has also been highlighted that the dentate gyrus plays a critical function in mediating processes such as recall of sequential information and short-term memory (Kesner, 2007). Subfields analysis related to AD has found that larger dentate gyrus volume is p redictive to the better follow-up cognitive and memory performance, emphasizing the key and neuroprotective role of dentate gyrus in preserving the memory function (Broadhouse et al., 2019). Fimbria is a small bundle of white matter fibers along hippocampal superior surface, and severs as a structural bridge between the hippocampus and rest of the brain (Au et al., 2021). It has been proposed that the integrity of fimbria is important for the preservation of hippocampus function in memory. The subiculum projects to the amygdala, entorhinal regions, mamillary nuclei and anterior and middle thalamic nuclei, and furtherly to frontal areas (Braak et al., 1996). Subiculum has been consistently reported to be affected by AD (Apostolova et al., 2006; Apostolova, Mosconi, et al., 2010; Apostolova, Thompson, et al., 2010). Previous studies have suggested that the prevalent atrophy of the presubicular-subicular complex is apparent in the early phase of AD and presubiculum and subiculum could be considered as possible predictors of verbal memory performance in MCI (Carlesimo et al., 2015). The hippocampal fissure is the inferolateral extension of the transverse fissure between the dentate gyrus and the subiculum (Bronen & Cheung, 1991; Duvernoy et al., 2005). Widen hippocampal fissure has been regarded as an indicator of AD (Li et al., 2018). Our results suggested that the integrity of input pathway (from molecular layer to core areas as dentate gyrus and CA4) and output pathway (through fimbria and presubicular-subicular pathway to thalamus and the rest of the brain) were affected by AD pathology.

Significantly, our results showed that the left subiculum, left molecular layer, left dentate gyrus and left CA4 were positively associated with VSTM accuracy or capacity, and the bilateral fimbria were negatively correlated with RT of VSTM in the control group, while no significant association was found in the aMCI group. the current results might reflect the deterioration of integrity of hippocampus for information input or output in response to the VSTM deficits in aMCI. As the hub for memory-related processes, hippocampal deterioration would disrupt the intricate neural networks responsible for memory encoding and retrieval. VSTM is not solely reliant on the hippocampus; it involves a complex interplay of brain regions and networks. These networks include the prefrontal cortex, parietal cortex, and regions within the occipital lobe (Prabhakaran et al., 2000; Ranganath et al., 2004; Rissman et al., 2008; von Allmen et al., 2014). Previous study has suggested the thalamus-hippocampus-prefrontal mechanism supported the memory (Aggleton & O’Mara, 2022). The interaction between thalamus and hippocampus has been suggested to support short-term memory (Arrieta-Cruz et al., 2010; Chengyang et al., 2017). In individuals with aMCI, disruptions in these networks may contribute to VSTM deficits, highlighting the need for a comprehensive understanding of the neural correlates of VSTM in this population. The current results highlighted the critical role of this thalamus-hippocampus-prefrontal mechanism in the AD pathology.

## Conclusion

The present study suggested that hippocampal atrophy, especially in subfields related to information input and output (e.g., molecular layer, DG, subiculum), may have contributed to VSTM impairment in aMCI by disrupting hippocampal-cortical communications. The pathway between hippocampus-thalamus-frontal cortical was highlighted. This finding adds to increasing evidence of hippocampal engagement in short-term memory processes and points to VSTM impairment as a potential neuropsychological indicator for MCI and AD.

## Notes

### Competing Interest Statement

The authors have declared no competing interest.

### Summary of Updates

Introduction and discussion has been updated

