## Supplementary Table for "Precise Alterations in Hippocampal Neural Deterioration and Neural Correlates of Visual Short-Term Memory in Individuals with Amnestic Mild Cognitive Impairment"

Supplementary Table 1. demographic characteristics.

|  | aMCI  (n = 55) | NC  (n = 68) | Between-group differences |
| --- | --- | --- | --- |
| Age (years) | 72.42 (6.69) | 70.31 (6.19) | t(121) = 1.81, *p* = 0.072^a^ |
| Gender (male: female) | 23:32 | 44:24 | $\chi^{2}$ = 6.42, *p* = 0.011^b^ |
| Education (years) | 11.33 (2.65) | 12.43 (2.79) | t(121) = -2.22, *p* = 0.028^a^ |
| MMSE | 24.86 (2.48) | 27.47 (1.65) | W = 638.5, *p* < 0.001^c^ |
| MoCA | 17.07 (3.34) | 24.24 (2.48) | t(121) = -13.64, *p* < 0.001^a^ |

Values are mean (standard deviation). Abbreviation: aMCI, amnestic mild cognitive impairment; NC, normal controls; MMSE, Mini-Mental State Examination; MoCA, Montreal Cognitive Assessment.

^a^ independent two-sample t-test, ^b^ chi-square test, ^c^ Mann-Whitney U test.
